# Dietary Advanced Glycation End-Product Consumption Leads to Mechanical Stiffening of Murine Intervertebral Discs

**DOI:** 10.1101/342691

**Authors:** Divya Krishnamoorthy, Robert C. Hoy, Devorah M Natelson, Olivia M. Torre, Damien M Laudier, James C. Iatridis, Svenja Illien-Jünger

## Abstract

Back pain is a leading cause of disability strongly associated with intervertebral disc (IVD) degeneration. Reducing structural disruption and catabolism in IVD degeneration remains an important clinical challenge. Pro-oxidant and structure-modifying advanced glycation end-products (AGEs) contribute to obesity and diabetes, which are associated with increased back pain, and accumulate in tissues due to hyperglycemia or ingestion of foods processed at high heat. Collagen-rich IVDs are particularly susceptible to AGE accumulation due to their slow metabolic rates yet it is unclear if dietary AGEs can cross the endplates to accumulate in IVDs. We apply a dietary mouse model to test the hypothesis that chronic consumption of high AGE diets results in sex-specific IVD structural disruption and functional changes. High AGE diet resulted in AGE accumulation in IVDs and increased IVD compressive stiffness, torque range, and failure torque, particularly for females. These biomechanical changes were likely caused by significantly increased AGE crosslinking in the annulus fibrosus, measured by multiphoton imaging. Increased collagen damage measured with collagen hybridizing peptide may be a risk factor for IVD degeneration as these animals age. The greater influence of high AGE diet on females is an important area of future investigation that may involve AGE receptors, known to interact with estrogen. We conclude high AGE diets can be a source for IVD crosslinking and collagen damage known to be important in IVD degeneration. This suggests dietary and other interventions that modify AGEs warrant further investigation and may be particularly important for diabetics where AGEs accumulate more rapidly.

**Summary Statement:** Dietary advanced glycation end-products (AGE) lead to sex-specific intervertebral disc structural and functional changes and may be targeted for promoting spinal health especially in diabetes where AGEs form rapidly.

## INTRODUCTION

Intervertebral disc (IVD) degeneration is a progressive condition and major contributor to back pain, which is a leading cause of global disability and absence from work (Hartvigsen et al., 2018, Adams and Roughley, 2006). Pain from IVD degeneration arises from structural disruption, catabolism, and chronic inflammatory conditions known to accumulate in IVD damage and degeneration. Consequently, it is a clinical priority to identify ways to reduce IVD damage. Risk factors for IVD degeneration are multifactorial, and recent studies have shown that Diabetes Mellitus (DM), obesity or being overweight add significant morbidity to back pain and IVD degeneration (Liuke et al., 2005, Takatalo et al., 2013, Jakoi et al., 2017, Samartzis et al., 2013, Fabiane et al., 2016) as they increase the risk for herniation (Sakellaridis, 2006), spinal stenosis (Anekstein et al., 2010) and spinal surgery complications (Guzman et al., 2014). DM and obesity are increasing at alarming rates, making this a growing high-risk population. Furthermore, DM and obesity contribute to chronic inflammation, catabolism, and altered biomechanics on the spine (Fields et al., 2015, Hillson, 2018, Illien-Junger et al., 2013), and we believe understanding risk factors for DM and obesity will also help elucidate novel risk factors for spinal diseases in the general population. Interestingly, sex differences in obesity and DM prevalence exist with DM women having increased risks for cardiovascular disease, myocardial infarction, and stroke mortality compared to DM men (Kautzky-Willer et al., 2016). The existence of sex-dimorphic pathologies in DM and other diseases motivates further investigation of sex-differences in spinal diseases.

Advanced glycation end-product (AGE) accumulation is a source for DM complications, and known to increase the risk for arthrosclerosis (Saremi et al., 2017) retinopathy and renal failure (Beisswenger et al., 1995). AGEs also accumulate to a greater degree in diabetic human IVDs where they are associated with increased matrix degrading enzymes (Tsai et al., 2014). In rats, type II DM was associated with degenerative changes in IVDs including glycosaminoglycan loss and IVD stiffening, which were again attributed to AGE accumulation (Fields et al., 2015). We previously demonstrated type I DM mice had increased IVD structural disruption and pro-inflammatory cytokines that were associated with AGE accumulation (Illien-Junger et al., 2013). We believe AGEs are a likely source of crosslinking and catabolism in the IVD, yet it is not clear if AGEs can accumulate in the avascular IVD from dietary ingestion or require hyperglycemia conditions from DM.

AGEs are highly oxidant compounds that can accumulate in tissues through endogenous (i.e. hyperglycemia) and exogenous (i.e. thermally processed foods) sources; 10% of dietary AGEs are absorbed via the intestine and released into the bloodstream (Koschinsky et al., 1997). Alarmingly, over the past 20 years, consumption of western diets consisting of highly processed foods has significantly increased, increasing the prevalence of obesity and DM (Cordain et al., 2005).

Recently, we showed in ageing pre-diabetic mice, that chronic ingestion of diets enriched with the specific AGE precursor (methylglyoxal) accelerated age-related vertebral bone loss and induced AGE accumulation within the endplate (Illien-Junger et al., 2015). When further investigated, ingestion of dietary AGEs induced sex dependent bone loss with inferior biomechanical properties in young (6m) female mice (Illien-Junger et al., 2018). These studies provided the first evidence that dietary AGEs had a direct effect on vertebral structure and function, and also showed that this effect was sex-dependent. Furthermore, immunostaining showed strong AGEs in the endplate and it remains unclear if dietary AGEs can accumulate in IVDs and contribute to structural or catabolic changes in the IVD known to be present in degeneration. Therefore, the aim of the current study was to assess the effects of dietary AGEs on structure and function of IVDs in female and male mice. We hypothesize that chronic ingestion of a high AGE diet will accumulate in IVDs and result in sex-specific IVD structural disruption and functional changes involving increased AGE crosslinking and collagen damage.

## RESULTS

### Dietary AGEs led to AGE accumulation in female IVDs

H-AGE diet led to significant accumulation of AGEs in IVDs (p=0.003, Figure 2) of female H-AGE compared to female L-AGE mice as seen by western blot analysis. This is in line with the observed increase in circulating AGEs found in H-AGE compared to L-AGE females as seen through serum ELISA analysis (16.9±4.3 U/mL vs. 9.5±4.8 U/mL, p=0.01). Interestingly, body weights (26.6±4.1 g vs. 26.3±2.6 g, n.s.) and fasting blood glucose (79.2±15.3 mg/dL vs. 84.2±18.7 mg/dL, n.s.) levels were not different between H-AGE and L-AGE females. No differences in AGE levels in IVDs or serum were observed for male mice (Serum AGE male: 11.7 ±4.2 U/mL vs. 8.3±.1.6 U/mL, n.s.). While male H-AGE mice had slightly decreased body weight and increased fasting blood glucose compared to L-AGE mice (body weight 27.3±1.3 g vs. 29.6±1.2 g, p=0.002; glucose: 88.4±11.6 mg/dL vs. 71.2±10.6 mg/dL, p=0.006), fasting blood glucose levels of all mice were below prediabetic levels (<100mg/dL) and body weights were also in normal ranges of this strain (Jackson_Laboratory). These data show that dietary AGEs accumulate systemically and can accumulate within IVD tissue in the absence of diabetes or obesity in female mice, while no differences were observed in male mice.

**Fig. 1.**
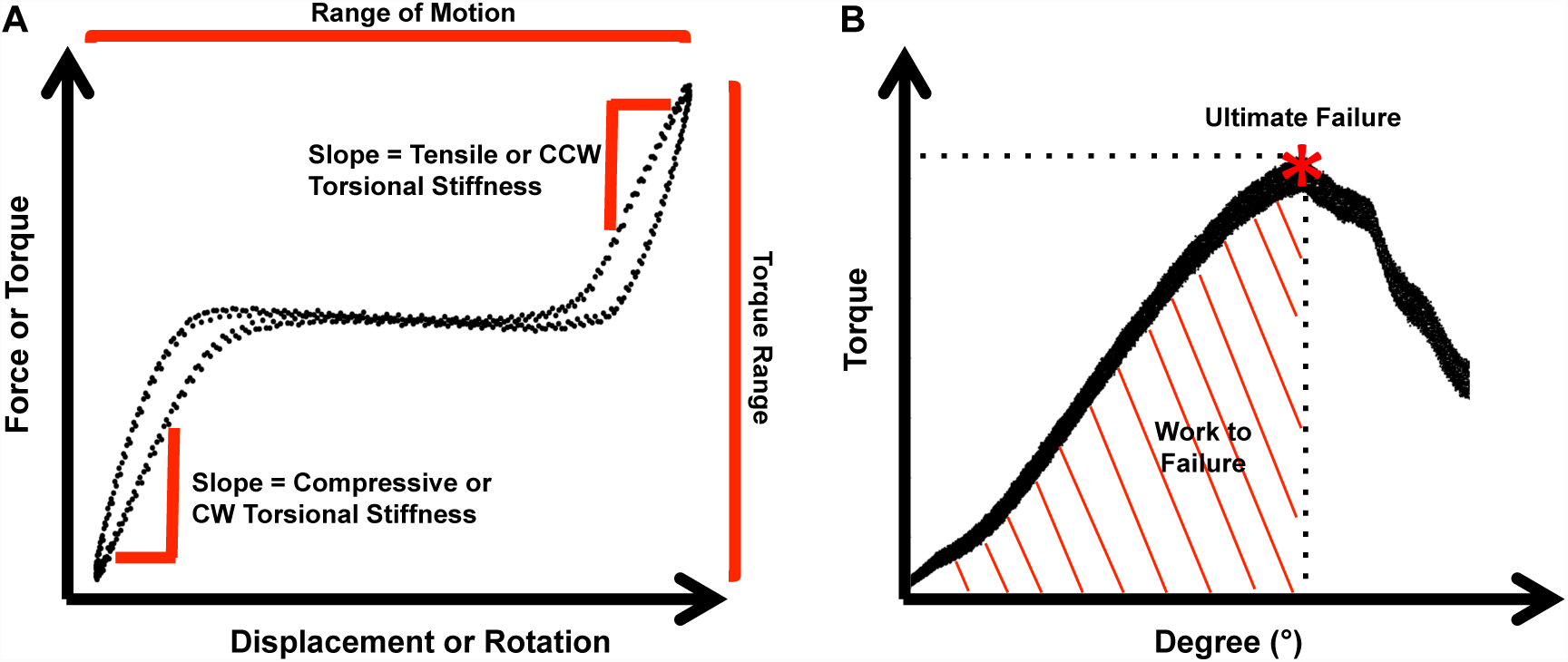
Biomechanical testing analyses. (A) Schematic of force-displacement curves for axial tension-compression and torsional testing showing linear regions used for stiffness measurements, regions measured for axial range of motion and Torque range. (B) Torsion to failure curve schematic showing point of ultimate failure and area under curve calculated as work done to failure.

**Fig. 2.**
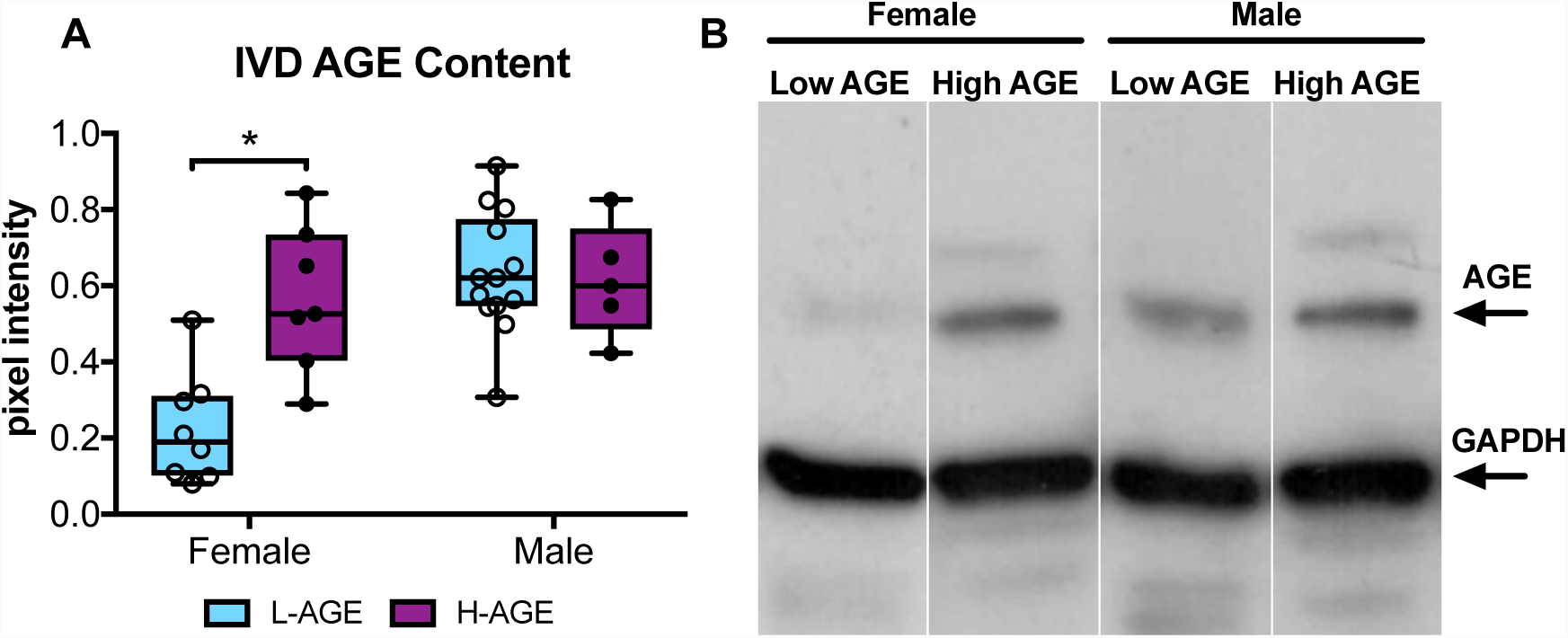
AGE accumulation in the IVD. (A) Western blot analysis showed greater AGE protein content in H-AGE female (n= 7) IVDs compared to L-AGE females (n=9), with no differences between H-AGE (n=5) and L-AGE (n=13) males. (B) Representative AGE western blot with representative bands for AGE and GAPDH, (internal control). Due to low protein concentration within some specimens, some samples needed to be pooled to warrant correct measurements. Data are presented as box plots from min to max ± s.d. P values are based on two-tailed unpaired Student’s t-test with Bonferroni correction and significant if p ≤ 0.05.

### Dietary AGEs led to increased axial compressive stiffness of female IVDs

Functional assessment of IVDs was performed by axial compression-tension and torsional biomechanical testing on coccygeal IVD motion segments. IVDs of female H-AGE mice had significantly increased compressive stiffness (p<0.001, Figure 3A) and increased torque range (p=0.031, Figure 3E) compared to female L-AGE mice. Failure analysis indicated that ultimate failure occurred at a greater torque in female H-AGE compared to L-AGE mice (p=0.05, Figure 3F) and similarly, work done to failure, which is comparable to material toughness, was also increased in female H-AGE compared to L-AGE mice (p=0.04, Figure 3H). No differences were observed for tensile stiffness, axial range of motion or torsional stiffness. Male IVDs were not affected by diet in any of the measured biomechanical parameters. These data show that motion segment behavior in compression and torsion is altered in female mice on a H-AGE diet and potentially indicates stiffening of annulus fibrosus fibers in the IVDs.

**Fig. 3.**
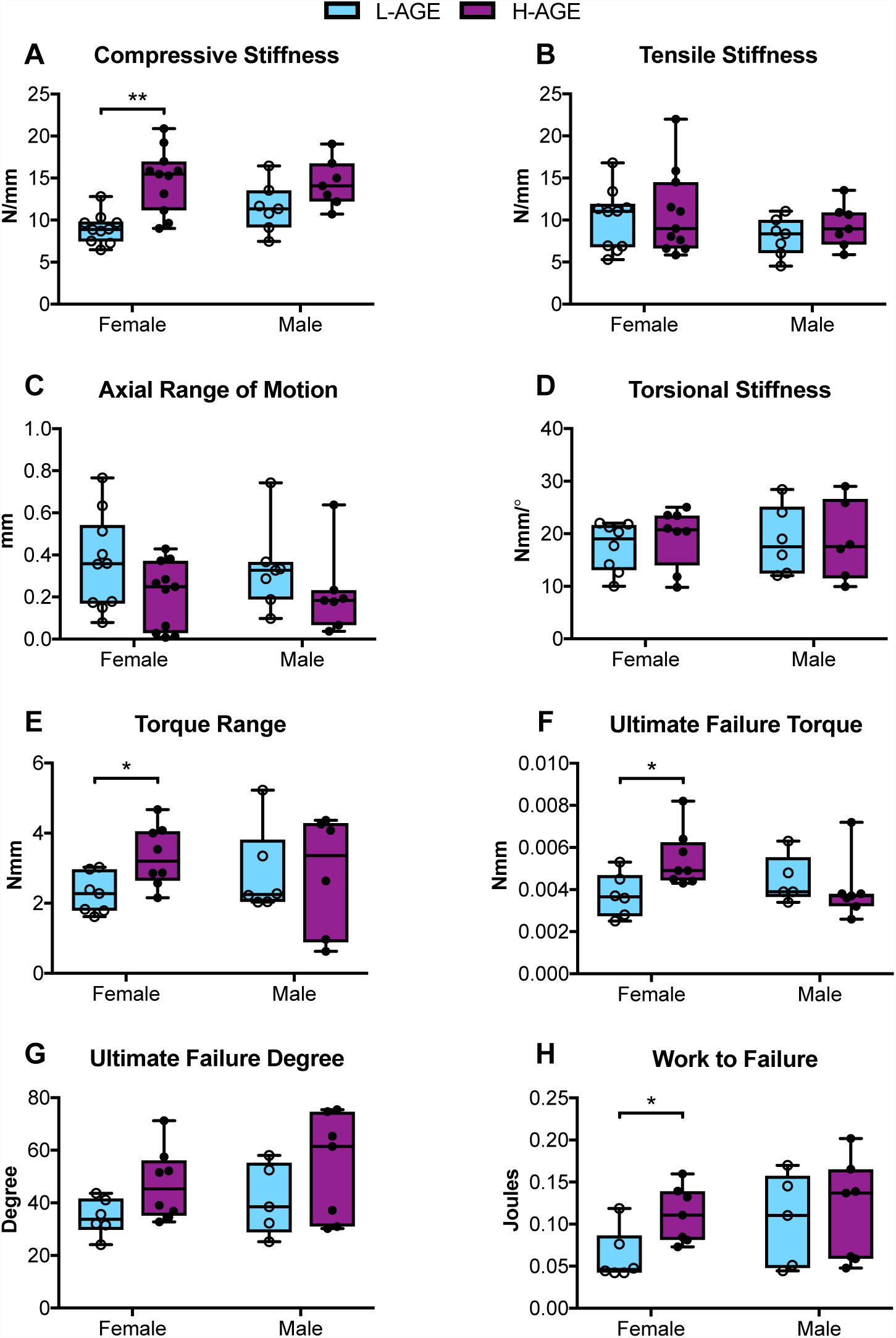
Tension-compression and torsion biomechanical testing. (A) Compressive stiffness increased in H-AGE F (n= 11) compared to L-AGE F (n=10), with no differences in (B) tensile stiffness or (C) axial range of motion or (D) torsional stiffness. (E) Torque range was increased in H-AGE F (n=8) compared to L-AGE F (n=7) Failure analysis revealed ultimate (F) failure torque and (H) work to failure increased in H-AGE F (n=7) compared to L-AGE F (n=6) with no difference in the (G) ultimate failure degree. No differences were detected between L-AGE M (n=5) and H-AGE M (n=7). Data are presented as box plots from min to max ± s.d. P values are based on two-tailed unpaired Student’s t-test with Bonferroni correction and significant if p ≤ 0.05.

### Dietary AGEs induced changes in annulus fibrosus organization of female IVDs

To assess morphological changes in the IVD, PR/AB stained sections were imaged under DIC and polarized filters. DIC imaging revealed no major changes in morphology in the NP or endplate (Figure 4A). However, when imaged under polarized filters, AF collagen fibers of female mice appeared brighter in the HAGE compared to L-AGE group, with no differences between male IVDs (Figure 4B). These differences were most prominently seen in the anterior AF where the collagen lamellae appear as a green/yellow color. These data indicate a difference in birefringence of collagen fibers in the H-AGE and L-AGE females.

**Fig. 4.**
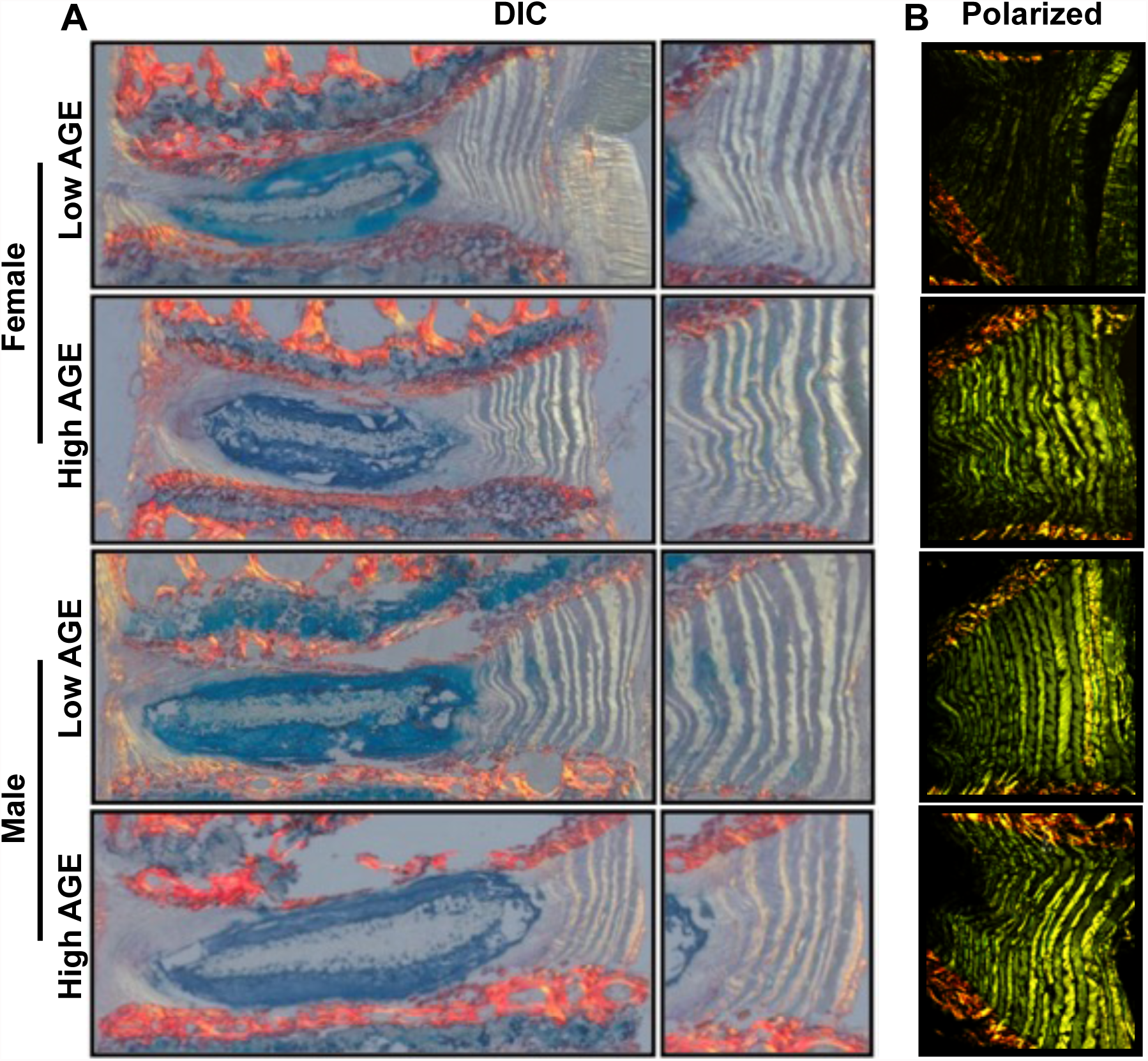
Histological analysis of IVD morphology. Representative sagittal sections stained with picrosirius red/alcian blue imaged under (A) differential interference contrast (DIC) and (B) polarized light. No differences were observed in end plate and nucleus pulposus between females of either diet. Polarized images showed a difference in the brightness of AF collagen fibers between the L-AGE (n= 5) and H-AGE (n=8) females. No differences were observed between L-AGE (n= 5) and H-AGE (n=7) males.

### Dietary AGEs impaired collagen quality in female IVDs

To investigate collagen quality we used two-photon imaging to quantify collagen SHG intensity and a collagen hybridizing peptide that reveals damaged collagen molecules. In female, but not in male mice, SHG measured in the anterior AF appeared less bright in H-AGE compared to L-AGE photomicrographs (Figure 5A). This was confirmed with quantitative measurements of the SHG intensity, which was significantly reduced in female H-AGE compared to L-AGE IVDs (p=0.037, Figure 5C). Furthermore, in the same region, measurement of TPEF/SHG, or total AGE auto fluorescence (Marturano et al., 2014) was increased in female H-AGE compared to L-AGE IVDs (p=0.03, Figure 5B). CHP staining of the same histological sections revealed significantly increased CHP+ green fluorescence staining in the AF of female H-AGE compared to L-AGE IVDs (p=0.03, Figure 6), indicating greater collagen damage. Alignment of collagen fibers (measured through fiber coherency) or lamellae organization (measured through a kinking factor) appeared to be not affected by AGE accumulation. No differences for any parameter were observed in male mice. These data indicate that H-AGE diet led to AGE accumulation and collagen damage in the AF of female mice only.

**Fig. 5.**
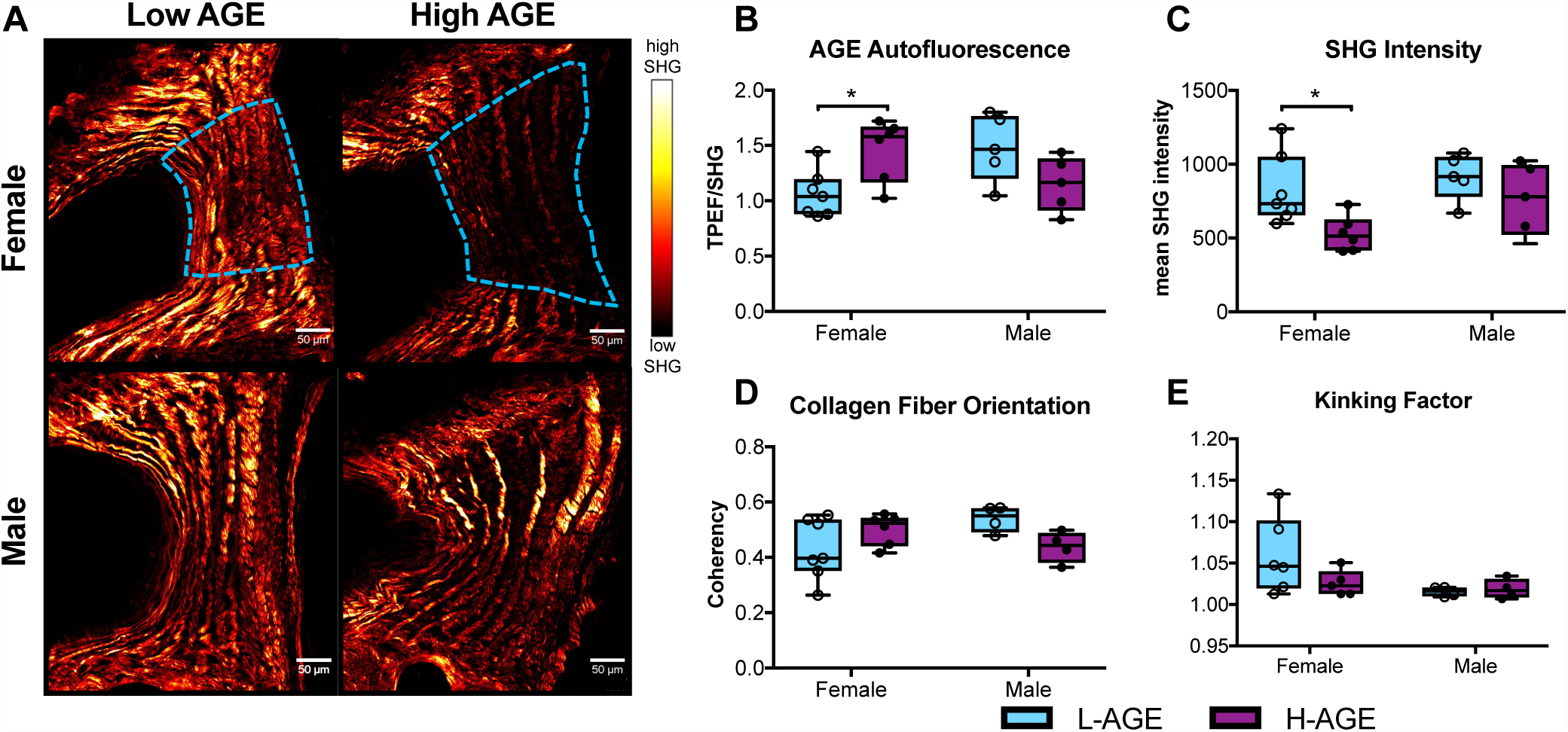
SHG of collagen in annulus fibrosus tissue. (A) Representative images with region of interest outlined in blue (B) AGE autofluorescence increased and (C) SHG intensity was reduced in H-AGE F (n=6) compared to L-AGE F (n=7) with no differences in (D) collagen fiber organization or (E) kinking of the lamellae. No differences were detected between males (L-AGE M and H-AGE M, n=5) Data are presented as box plots from min to max ± s.d. P values are based on two-tailed unpaired Student’s t-test with Bonferroni correction and significant if p ≤ 0.05.

**Fig. 6.**
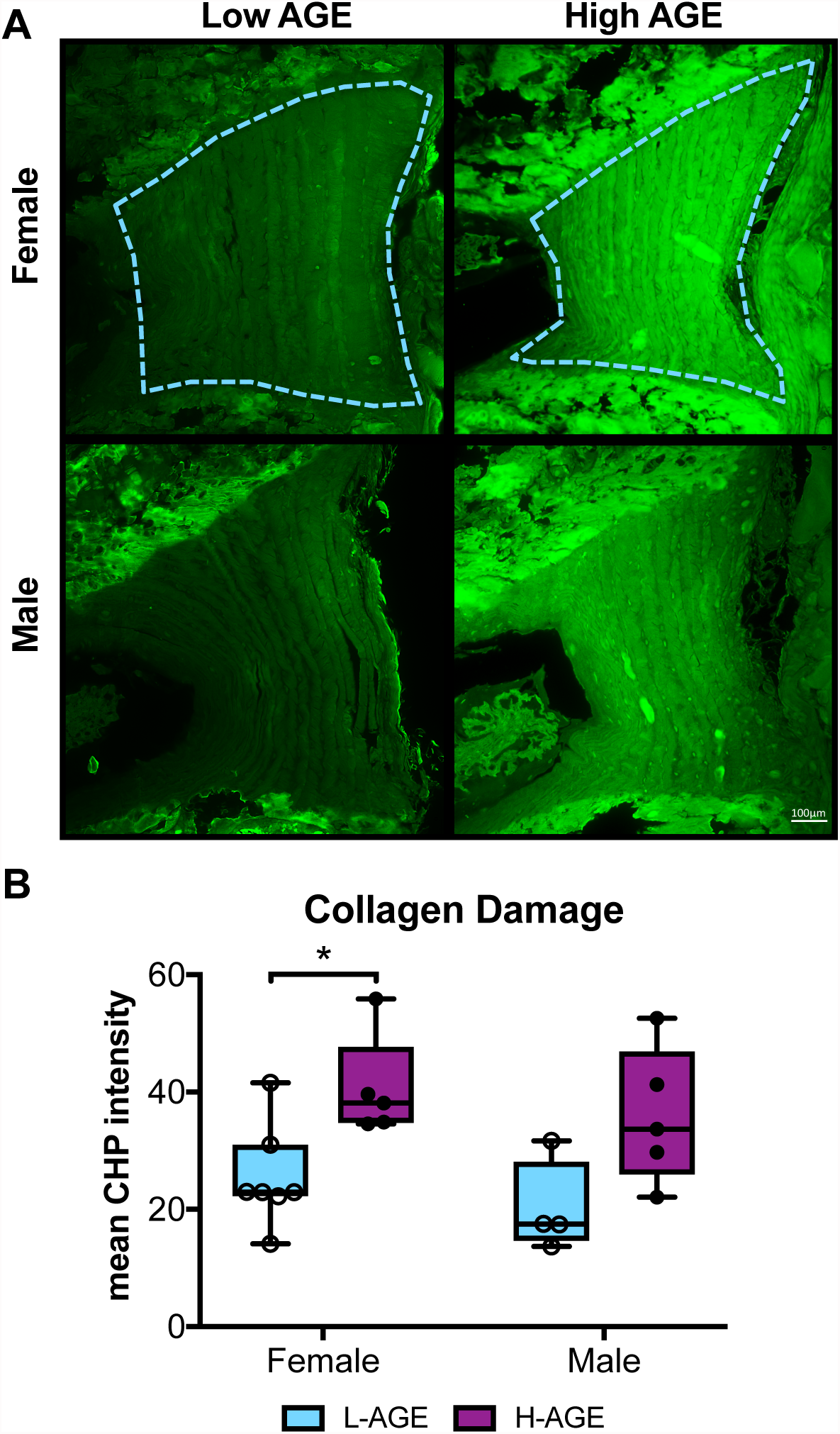
Molecular assessment of collagen. (A) CHP stained fluorescent images of anterior AF exhibit increased CHP staining in H-AGE compared to L-AGE in females and males. (B) Quantification of CHP mean intensity shows significant collagen damage in H-AGE F (n= 5) compared to L-AGE F (n=7). No differences were detected in between males (L-AGE M n=4, H-AGE M n=5). Data are presented as box plots from min to max ± s.d. P values are based on two-tailed unpaired Student’s t-test with Bonferroni correction and significant if p ≤ 0.05.

## DISCUSSION

It is a priority to identify sources for structural disruption and matrix damage in IVD degeneration since these conditions are known causes for painful conditions. AGE accumulation in tissues is a natural function of aging but is accelerated in conditions such as DM and can be increased by consumption of poor diets (Tessier et al., 2016, Koschinsky et al., 1997). This study demonstrated that, especially for females, consumption of processed diets that are high in AGE, resulted in IVD AGE accumulation causing functional changes including increased compressive stiffness, torque range and torque failure properties. Significant alterations in collagen quality were also observed and the significantly increased AGE crosslinking was most likely responsible for the observed biomechanical changes in these young animals although significantly increased collagen damage may have also contributed and will likely result in increase failure risk as IVDs age. These effects of high AGE diet on IVDs were sex-dependent, since only female mice were significantly affected and no significant effects were detected for males, although multiple parameters had means following similar trends as females. The absence of hyperglycemia and obesity suggests that the majority of AGEs measured in the IVDs and subsequent functional and structural changes in this study were a direct result of the dietary AGEs.

AGEs, including pentosidine, crosslink proteins to alter tissue structure and function, and can be a marker of protein aging and turnover (Sivan et al., 2006). The propensity to crosslink long living proteins such as aggrecan and collagen has been shown to increase stiffness and brittleness in tissues like articular cartilage (Verzijl et al., 2002), bone (Vashishth et al., 2001) and even the IVD (Wagner et al., 2006). AGE collagen crosslinks accumulate in IVDs with aging, being implicated with loss of IVD integrity and advancing the pathogenesis of the degenerative process (Pokharna and Phillips, 1998). Moreover, in vitro glycosylation of the AF specifically, increases tissue stiffness and brittleness (Wagner et al., 2006). As shown in this study, AGEs can also accumulate with poor diet to potentially accelerate aging. We also observed increased AF stiffness as detected with significantly increased torque range and torque failure properties in females. Our results also reflect other studies, which demonstrated increased compressive stiffness due to increased collagen crosslinking (Barbir et al., 2010). It is noteworthy that we did not observe differences in tensile stiffness during axial tension-compression testing since AF collagen fibers, if highly crosslinked, likely also resist tension when tensed axially. Yet, it is possible that crosslinked IVDs exhibit a greater increase in circumferential tension and axial compression compared to axial tension and radial compression (Chuang et al., 2007). Furthermore, we also identified collagen damage that may act in opposition to the increased crosslinking to reduce sensitivity of some of these structural biomechanical properties to specific collagen changes. In the context of the literature, the majority of biomechanical changes observed in this study are consistent with increased collagen crosslinking from AGE accumulation.

Collagen damage as shown through CHP binding, was significantly increased in the AF of female IVDs with high AGE diet, yet these results did not seem to have a detectable effect on IVD biomechanics. The increased collagen damage appears to contrast the significantly increased torsional failure strength and greater amount of work to reach torsional failure in the same group. CHP is a highly sensitive measure of collagen damage, which may not have accumulated sufficiently in these samples to result in biomechanical differences. The fiber-reinforced laminated composite structure of the AF has multiple levels of redundant fibers that are highly effective at maintaining function and resisting crack propagation (Iatridis and ap Gwynn, 2004). Additionally, the spine undergoes various motions that combine complex loading conditions that accumulate with aging, yet even damaged collagen fibers would require high magnitudes of loading and/or older aged animals to significantly degrade biomechanical function to detectable levels. We also evaluated caudal motion segments since they are easier to grip and test biomechanically than lumbar spinal motion segments in these small mouse tissues, and caudal IVDs are subjected to lower peak forces than spinal motion segments (Elliott and Sarver, 2004). While our data suggests that biomechanical changes are driven by AGE crosslinking in the AF, we also observed that these pro-oxidative AGEs lead to collagen damage, which is likely to increase risk of IVD injury at higher forces or with increased aging.

Histological assessment of the AF morphology revealed reduced SHG from the collagen fibers. Decreased SHG intensity can indicate increased crosslinking due to glycation (Tseng et al., 2010, Kim et al., 2000) and structural disorder of the collagen fibrillar array (Reiser et al., 2007). Glycation has been shown to disrupt collagen organization in multiple ways: AGEs can alter fibril organization through increased fibril packing and diameter (Bai et al., 1992, Hwang et al., 2011); AGE adducts can twist and distort the collagen fibrillar array (Kim et al., 2000); and AGEs can alter molecular packing by increasing the distances between collagen chains and molecules (Tanaka et al., 1988). Our data suggests that higher order alterations to fibril structure such as fibril diameter and packing due to AGE crosslinking are likely contributing to loss of collagen SHG seen in the H-AGE female IVDs, however, the assessment of fibril structure would require much higher resolution techniques.

This study demonstrated increased CHP staining, as a direct measure of damage to the triple helical structure of collagen molecules, in the H-AGE female group, which may also account for the reduced SHG intensity. Alterations in fibril packing can increase the susceptibility to enzymatic digestion (McBride et al., 1997) and collagen degradation, which in turn can contribute to the loss of collagen SHG intensity (Hwang et al., 2017). It has been proposed that AGE crosslinking can lead to mechanically induced breakdown of the collagen triple helical structure by causing forces, transmitted through the intermolecular AGE crosslinks, to accelerate enzyme degradation (Bourne et al., 2014). However, AGEs can also induce enzymatic activity and matrix catabolism, for example by binding to the receptor for AGEs (RAGE) (Ott et al., 2014). Some have suggested an important role for the AGE/RAGE pathway in diabetes and AGE induced IVD degeneration (Fields et al., 2015, Illien-Junger et al., 2016) however future studies will investigate the role of this interaction further especially as it pertains to dietary AGEs. Together with the literature, our assessment of SHG intensity and CHP binding suggests that dietary AGEs lead to increased glycation within the AF collagen matrix, inducing collagen fibrillar disorder and molecular damage.

Causes for the sex-dependent effects of dietary AGEs warrant future investigation, especially because estrogen is known to regulate cell metabolism (Mauvais-Jarvis et al., 2013) and to influence the receptor for AGEs (Tanaka et al., 2000). We previously have shown that dietary AGEs affect bone mechanical properties mostly in young female mice (Illien-Junger et al., 2018), which further emphasizes the importance of assessing male and females separately. Furthermore, the variations in vertebrae and IVDs also highlight the importance of evaluations of all spinal structures, which interact and may influence each other. While we did not control for the estrus cycle, which may have contributed to a greater variance within the female data, this did not seem to be important in our measurements since many significant differences were identified for female but not male IVDs. Lastly, while some studies have shown AGE accumulation in the nucleus pulposus, having effects on IVD hydration and pressurization (Fields et al., 2015, Illien-Junger et al., 2013), we did not observe any alterations to NP morphology with histology, suggesting little contribution of NP to the observed mechanical alterations and that changes to the AF are largely driving the functional changes in the female IVDs.

In conclusion, high AGE diets can be a source for IVD crosslinking and collagen damage that are known to be important in IVD degeneration and that can result in changes to functional IVD biomechanical behaviors These dietary AGE effects on the spine were significant for females but not males highlighting the importance of sex-dependent effects on spinal tissues. Results suggest that dietary and other interventions that modify AGEs warrant further investigation for promoting spinal health. Due to the increased risk for AGE accumulation with hyperglycemia in DM, we further conclude that targeting AGEs may be particularly important in promoting spinal health in DM patients.

## METHODS

### Mouse Model

After weaning, 21 female and 23 male C57BL/6J mice were each assigned to two iso-caloric diet groups, receiving either a low AGE chow (L-AGE; containing 7.6 μg/mg AGE; Test Diet Low AGE 5053; WF Fisher & Son CO, NJ, USA), or high AGE chow (H-AGE; 40.9 μg/mg; NIH-31 open formula chow autoclaved for 30 minutes at 120°C) (Table 1). Experimental animals were bred from parent mice that also consumed either a L-AGE or H-AGE diet. Mice were group housed and had ad libitum access to food and water. Mice were euthanized at 6 months of age by cardiac puncture. Prior to sacrifice body weights and fasting blood glucose measurements (16-hour fast) were acquired to establish body habitus and diabetic status.

**Table 1:** Animal Model

| **C57BL/6J mice** | **Low AGE Diet (L-AGE)**<br>**7.6 µg/mg AGE** | **High AGE Diet (H-AGE)**<br>**40.9 µg/mg AGE** |
| --- | --- | --- |
| Female | n=10 | n=11 |
| Male | n=13 | n=10 |

**Table 2:** Primary Study Measurements.

| **TISSUE** | **TEST** | **MEASUREMENT** |
| --- | --- | --- |
| Blood | Serum ELISA | Blood glucose<br>AGE measurement |
| Caudal IVD | Western Blot | AGE measurement |
| Caudal IVD C4-C5 | Mechanical Testing | Axial, Torsional and Failure tests |
| Lumbar IVD L4-5 | Histology<br>(Picrosirius/Alcian blue) | DIC, polarized light & SHG imaging<br>collagen denaturation |

### Tissue Harvest

Following sacrifice, caudal (C) IVDs from C5 through C10 were carefully dissected from each tail and stored in -80°C until use for western blot analysis. For biomechanical testing, C4-C5 bone-IVD-bone motion segments were dissected from mouse-tails. Motion segments were prepared by making a parallel cut (transverse to the long axis of the spine) through the C vertebrae and included the C4-C5 IVD. The facet joints and all surrounding soft tissue were removed. Motion segments were wrapped in phosphate buffer saline (PBS) soaked paper towels and stored in -80°C until day of testing. Lumbar (L) L3-L4 and L4-L5 segments were dissected and fixed in 10% neutral buffered formalin prior to resin embedding for histological analysis.

### Protein measurements for AGE Quantification

For protein extraction, IVDs were placed in lysis buffer (T-PER tissue protein extraction reagent, Thermo Scientific, Waltham, MA, USA) + protease inhibitor (Protease inhibitor cocktail, Bimake, Houston, TX, USA) and sonicated. Following centrifugation, supernatants were collected and proteins were separated by sodium dodecyl sulphate polyacrylamide gel electrophoresis (SDS-PAGE; Bio-Rad Laboratories, Hercules, CA, USA). 20 µg protein samples were loaded on gradient gels (4–20 %; Bio-Rad Laboratories, Hercules, CA, USA). After electrophoresis, proteins were transferred on nitrocellulose blotting membranes (Protran BA Cellulosenitrat; Schleicher & Schuell, Dassel, Germany). Following the protein transfer, membranes were incubated for 30 minutes in blocking buffer (5 % milk powder in PBS + 0.2% Tween^®^ 20) and then incubated with the primary antibodies, AGE (1:3,000; ab23722; abcam, Cambridge, MA, USA) and GAPDH (internal control; 1:20,000; ab181602; abcam, Cambridge, MA, USA), overnight at 4°C. The next day the membranes were washed in washing buffer (PBS + 0.2% Tween^®^ 20) and incubated with an Eu-labeled secondary antibody (Molecular Devices, San Jose, CA, USA) for 1 hour at room temperature. After washing, the membranes were dried and scanned using the ScanLater (Molecular Devices, San Jose, CA, USA) Western Blot cartridge in a SpectraMax i3 system (Molecular Devices, San Jose, CA, USA). Membranes were analyzed by ScanLater software.

### Biomechanical Testing

Prior to mechanical testing, motion segments were thawed and hydrated in cold 1X PBS. Axial and torsional testing was performed on the ElectroForce 3200 and the AR2000x Rheometer (TA instruments, New Castle, DE, USA), respectively. For axial testing, each motion segment was placed in a 1X PBS bath at room temperature between custom designed fixtures and microvises. The test consisted of 20 cycles of tension and compression with peak loads of 0.5 N applied at a 0.1 mm/sec displacement rate (Elliott and Sarver, 2004). To recover IVD height after axial testing, motion segments were placed into in a 1X PBS bath for 5 minutes prior to torsional testing. Following an axial preload for 5 min at 0.1 MPa axial stress, torsional testing was performed and consisted of 20 cycles of rotation to ±10° at a frequency of 1 HZ followed by a 1°/sec continuous rotation to failure (Michalek et al., 2010). Sample sizes were initially balanced yet final numbers vary between groups because some motion segments failed during tension-compression or torsion testing and thus were excluded from analysis. A custom written MATLAB code was used for calculating compressive stiffness, tensile stiffness, axial range of motion, torsional stiffness and toque range using the second to last cycle of every test. Stiffness measurements were defined as the slope of 20% of the top and bottom (linear region) of the force-displacement curves. Torsional stiffness is reported as the average of clockwise and counter-clockwise measurements. Ultimate failure torque and degree were calculated manually from the force-displacement curves and the work done to failure was calculated as the area under this curve.

### Histological Assessment

#### IVD Morphology

Fixed, calcified lumbar segments were embedded in poly(methyl methacrylate) (PMMA) and 5 µm sagittal sections were cut. Mid-sagittal sections were stained with Picrosirius Red and Alcian Blue (PR/AB) to assess collagen and proteoglycan content, respectively. PR/AB stained sections were then imaged under differential interference contrast (DIC) using the Axio Imager Z1 (Zeiss, Oberkochen, Germany) to assess overall IVD morphology and composition. The same samples were then imaged under polarized light using the LEICA DM6 B (Leica Microsystems, Illinois, USA) for qualitative assessment of annulus fibrosus (AF) collagen organization and arrangement. For all histological analysis, while sample sizes were initially balanced across groups, final numbers vary in each group because only mid sagittal sections were used for comparison and any non-sagittal sections were excluded from analysis, also any outliers due to technical errors with staining were also excluded.

#### Molecular Assessment of Collagen

PMMA embedded mid-sagittal lumbar sections were stained using Collagen Hybridizing Peptide-biotin conjugate (B-CHP; BIO300, 3Helix Inc; Salt Lake City, UT, USA) to detect the presence of collagen damage as described by (Hwang et al., 2017, Zitnay et al., 2017). This procedure includes the creation of a 2 μM CHP solution, heating of the solution at 80°C for 5 minutes to monomerize the peptides, quenching in an ice bath for 30 seconds, and application of the solution to the tissue. The tissue was incubated overnight, and then positive binding was detected using GFP labeled streptavidin (Dylight 488 Strepavidin; Vector Laboratories Inc, Burlingame, CA, USA).

### Second Harmonic Generation (SHG) Imaging

Unstained resin-embedded sections were imaged with an Olympus FV1000 MPE laser-scanning microscope (Olympus Corporation, Tokyo, Japan) at the Icahn School of Medicine at Mount Sinai Microscopy Core. Two-Photon excitation was done with the tunable Coherent Chameleon Vision II laser. Backward signal propagation was collected using the dedicated Olympus WLPLN water immersive 25X objective with a numerical aperture of 1.05. A stepwise approach was used to capture the second harmonic generation (SHG) signal of fibrillary collagen, and the Two Photon Excitation Fluorescence (TPEF) signal of total AGE auto fluorescence (Tang et al., 2007, Vashishth, 2009) in the anterior AF of each sample. Excitation for backward SHG (B-SHG) was performed at 910nm and recorded by a photomultiplier tube (PMT) at 440 +/- 10nm. The laser was then tuned to 740nm and the TPEF was recorded with the same detector at 440 +/- 10mm. All parameters (i.e. Laser Intensity, Gain, HV, and Dwell Time, Aspect Ratio) were held constant for the SHG and the TPEF imaging to allow comparison of image intensity. Optical slices were taken through the entirety of every section at a step size of 1.5 μm and a maximum intensity z-projection was performed for both SHG and TPEF. The B-SHG and TPEF intensities were assessed by measuring the average pixel intensity/area of the anterior AF using ImageJ.

### Fiber Orientation Analysis

Collagen orientation was quantified from the B-SHG images using OrientationJ (Rezakhaniha et al., 2012). To optimize recognition of AF fibers, the color balance was manually adjusted to maximize the observer’s ability to distinguish fibers. A Laplacian of Gaussian prefilter (σ = 1.0) was then applied to decrease noise and enhance the edge detection of the fibers. Four adjacent lamellae were selected from each anterior AF based on the following criteria; adjacent lamellae must have alternating dominate directions, the lamellae must span the height of the disc, and extreme changes in a dominant direction signified an interlammelaer boundary. Each lamellae was outlined as a region of interest and the coherency was measured. The coherency indicates if the local fibers within the region of interest are orientated in one direction; a coherency of 1 describes fibers that are in the same orientation, while a coherency of 0 describes fibers that are isotropic in the local neighborhood. The average coherency from the four regions of interest was reported.

### Statistics

All data are presented as box plots with minimum, first quartile, median, third quartile and maximum values with error bars representing the standard deviation. Two-tailed unpaired Student’s t-test with Bonferroni correction was applied between L-AGE F and H-AGE F and between L-AGE M and H-AGE M. All data were considered significant if p ≤ 0.05 and a trend if p ≤ 0.10.

## ACKNOWLEDGEMENTS

Multiphoton microscopy was performed in the Microscopy CoRE at the Icahn School of Medicine at Mount Sinai, supported with funding from National Institute of Health Shared Instrumentation Grant (1S10RR0266390). We thank Philip Nasser, Warren Hom and Dr. Rose Long for their assistance with biomechanics testing and analysis and Madeline Smith for polarized light imaging.

## AUTHOR CONTRIBUTION

Authors’ roles: DK performed and analyzed experiments, wrote the manuscript and was involved in all aspects of the submission. RCH, OMT and DMN contributed to the experimental work, reviewing and editing of the manuscript. JCI and SIJ conceptualized and designed the study, contributed to analysis and interpretation, and critically revised and edited the manuscript.

